# Shorter season options through transplanted fodder beet

**DOI:** 10.1101/090738

**Authors:** EN Khaembah, S Maley, WR Nelson

## Abstract

Fodder beet has distinct benefits such as high yield potential, excellent feed quality (especially metabolisable energy), and suitability for cool temperate climates. Production area has recently increased dramatically in New Zealand, primarily for non-lactating cow feed during winter but increasingly for other animals and times of the year.

Currently, establishing fodder beet requires intensive land cultivation and precision sowing of pelleted seed. It is generally regarded as a difficult crop to grow successfully. Competition from early season weeds means that multiple herbicide applications are commonly applied. Delaying the sowing date, until soil temperatures have risen enough for germination, limits the flexibility of this crop within farm rotations.

Transplanting is a plant establishment technique common in both forestry and vegetable crops. It simplifies establishment and reduces the risk of poor establishment.

Here we demonstrate that transplanting of fodder beet can be conducted successfully with low variability observed within the transplanted crop. Individual root volume and dry matter content are similar, whether crops are precision-drilled or transplanted. Our results suggest that transplanting is a financially feasible option for fodder beet establishment.

## Introduction

In New Zealand, demand for locally grown animal feed is important for economic, animal welfare and environmental benefits (Malcolm et al. 2016). Economic analyses of dairy systems incorporating fodder beet (*Beta vulgaris* L.) have confirmed the high value obtainable from this crop, with significant per hectare profit advantages by incorporating a higher proportion of feed energy from fodder beet compared to grass pasture, or by substituting fodder beet for cereal or imported palm kernel feeds (James 2015; Riley 2015). Fodder beet already contributes 41% as much feed for dairy cows in New Zealand compared to forage brassicas, but using only 22% as much land (Dairy NZ 2016).

Fodder beet is a close relative of both sugar beet and the vegetables beetroot and chard. Precision sowing for optimal yield is difficult because of erratic and slow germination resulting in plant losses and weed competition (Scott and Maley 2010; Gibbs 2014). The sugar beet industry has had a long history of using transplanting establishment in short season temperate climates (Scott and Bremner 1966; Moraghan and Torkelson 1972; Brinkmann 1986). In warmer climates, precision drilling remains the method of choice as establishment costs and risk of poor establishment are lower. Previous reports of difficulty in growing sugar beet crops in New Zealand reinforce the observations of erratic establishment of fodder beet (McCormick and Thomsen 1980; Kemp et al. 1994). Further, irregular crop establishment leads to erratic individual plant size within the population (Gibbs 2015), leading to difficulty estimating whole crop yields and complicating break feeding estimates.

We have previously reported on preliminary trials comparing fodder beet established by precision drilling or transplanting (Khaembah and Nelson 2016). In a similar small plot trial we focus here on the variation in crop establishment and growth between precision-drill and transplant methods.

## Methods and Materials

The trial was conducted at the New Zealand Institute for Plant & Food Research Ltd, Lincoln farm (43.625°S, 172.467°E, 12 m above sea level), Canterbury, New Zealand. A commercial fodder beet cultivar (‘Rivage’, Agricom Ltd, Christchurch) was used in this study. Seedlings were glasshouse-raised in the 144 Transplant Systems cell trays and transplanted by hand on 11 November 2015 when they were 38 days old. Seedlings were transplanted in four 20 m rows spaced at 45 cm and intra-row spacing of 25 cm (90,000 plants/ha). On the same day, pelleted seeds were precision-drilled in eight rows (four on either side of the transplanted plots) using an air-seeder at the rate of 110,000 seeds/ha. Row spacing of precision-drilled plots was 0.5 m. Pre-sowing/transplanting fertiliser and topdressings were applied as per common agronomic recommendations (Chakwizira et al. 2014). Irrigation and herbicides were applied as required.

### Measurements

Canopy light interception was measured at 9 to 33 day intervals from 17 November 2015 to 02 June 2016 using a portable Sunfleck ceptometer (AccuPAR model PAR-80; Decagon, Pullman, WA, USA). Measurements were confined to ±1 hour of local solar noon (Vaesen et al. 2001) and were taken on sunny days only.

Samples for dry matter (DM) and plant population density determination were harvested three times: 27 January 2016, 10 March 2016 and 08 June 2016 (final harvest). At each sampling time, plants were lifted by hand from quadrats randomly placed within the two inner rows of transplanted/precision-drilled plots Plant population and fresh weight were determined in the field. Subsamples of two plants from each plot sample were washed to remove soil, re-weighed, cut into small pieces and oven-dried at 60°C until a constant weight to determine total DM mass. At the final harvest, the volume of storage roots (sometimes referred to as bulbs) of subsamples was determined before drying samples for DM determination. Volume was determined by immersing storage roots in a bucket full of water and weighing the displaced water. A 1:1 weight to volume ratio was assumed i.e. 1 litre of water = 1 kg of water.

### Economic analysis

A cost of establishment and gross margin comparison was determined on a per hectare basis for both methods (Table 1). Costs estimated for precision-drilled fodder beet production in the Hawke’s Bay (Matthew et al. 2011) were also included in the comparison. We assumed the same costs for ground preparation and fertiliser for both methods. The major difference in cost results from both the nursery phase of growing the seedlings and the cost of transplanting itself. We estimated the seedling cost at NZ$35/1000 (including seed cost) based on a range of estimates from commercial seedling nurseries. We had more difficulty estimating a price for transplanting as it is currently more common for vegetable farmers to operate their own transplanting equipment rather than using external contractors. Although our estimates ranged from $600–$1,200/ha, because of high uncertainty, we have used the higher estimate. On this basis, the precision-drilled crop had an establishment cost of $2,879/ha and the transplants about double this at $5,977/ha.

**Table 1:**
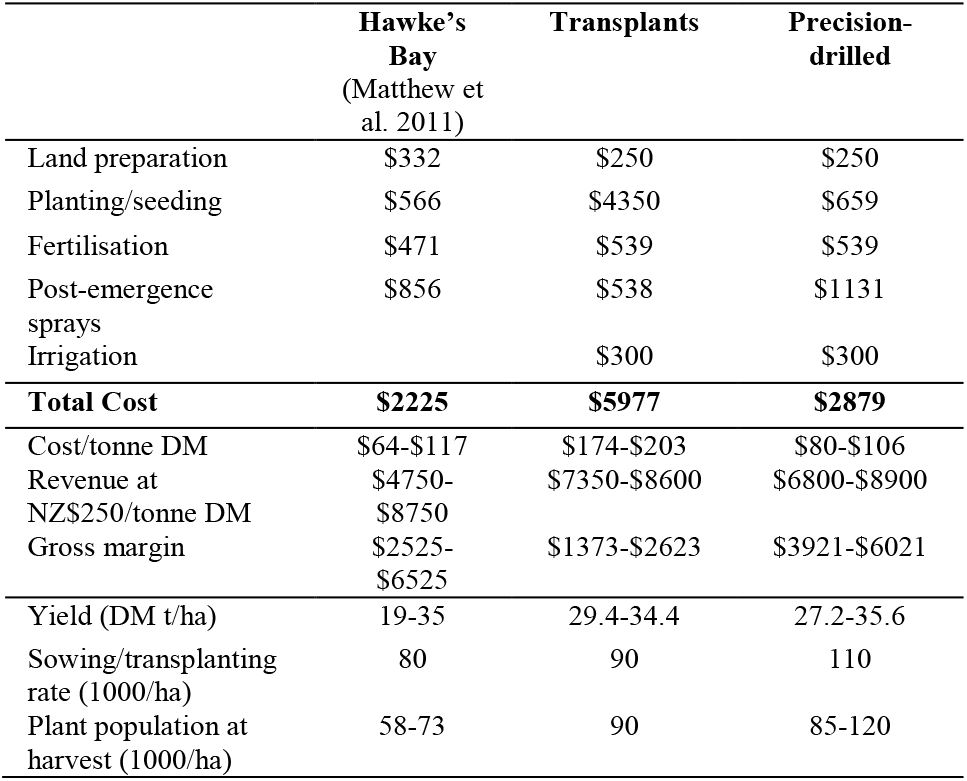
Cost estimates of plant establishment systems for fodder beet. Hawke’s Bay figures from three farms (Matthew et al. 2011). Prices in NZ$.

Gross margin for the crops was based on a current price of $250/tonne DM in the paddock. We have not estimated lifting, storage or feed-out costs as these crops are commonly grazed *in situ.*

## Results and Discussion

Establishment to canopy cover of the transplanted crop was again earlier than precision-drilled as reported previously (Khaembah and Nelson 2016). This gives very clear evidence for early season canopy cover advantage (Figure 1), reducing weed competition so that fewer herbicide applications were required (Table 1) and an early season light interception advantage (Figure 2). The transplants achieved 100% stand establishment at 90,000/ha, while the drilled crop averaged 105,200/ha (range = 85,000−120,000) calculated from sample quadrats (Figure 3).

**Figure 1:**
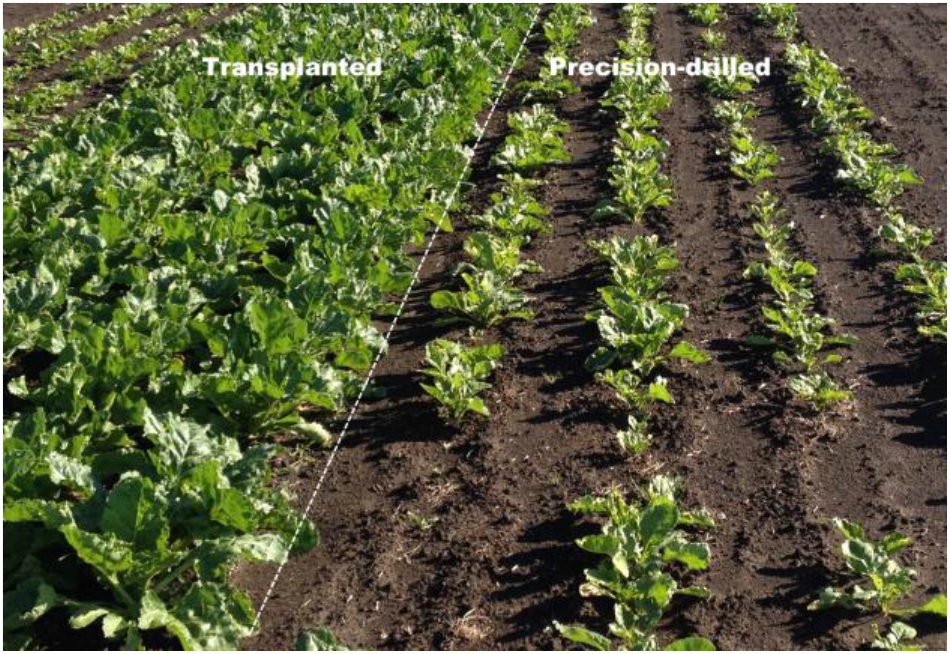
A bed of transplanted fodder beet showing near canopy cover compared to precision-drilled beds on either side at 4 weeks after sowing/transplanting. Note the variation in the drilled seedlings, both in plant population and plant size.

**Figure 2:**
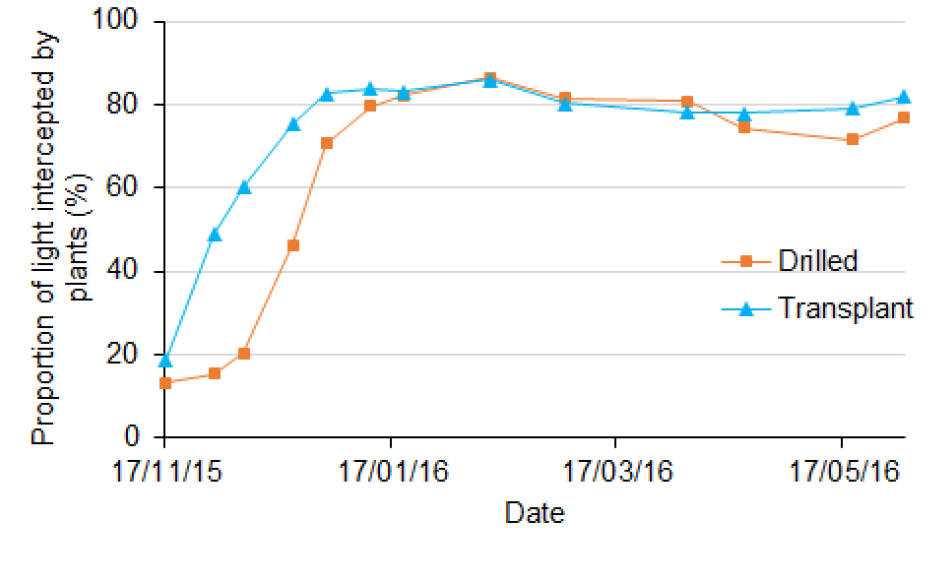
Proportion of light intercepted by plants showing the nearly two month advantage for transplants to reach full canopy cover.

**Figure 3:**
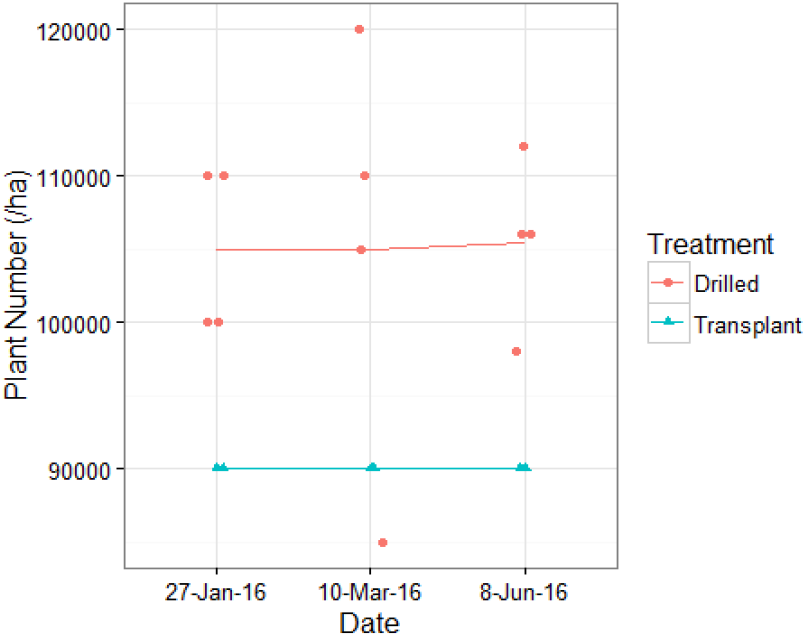
Plant populations of precision drilled and transplanted fodder beet, calculated from plant numbers within quadrats at three sampling periods through the growing season. The line is a simple average for each treatment and time point. The number of samples from plots at each harvest was two and four from the transplanted and drilled plots, respectively.

At the January harvest the transplanted plants achieved an early canopy cover evident by the higher yield, although this advantage over the precision-drilled crops appears to have reduced by the March harvest (Figure 4). This is also shown by the transplanted plants also exhibiting a higher %DM at this earlier stage (Figure 5). Total DM for both crops was essentially the same at approximately 31t DM/ha. This reflects a very high yield compared to commercial drilled crops commonly reported below 20t DM/ha (Scott and Maley 2010) and seldom as much as 25t DM/ha (Milne et al. 2014). However, a feature of commercial precision drilled crops is the variability within the crop as plants respond to large differences in plant population through areas of over sowing, areas of reduced population and occasional large patches far too big for neighbouring plants to compensate.

**Figure 4:**
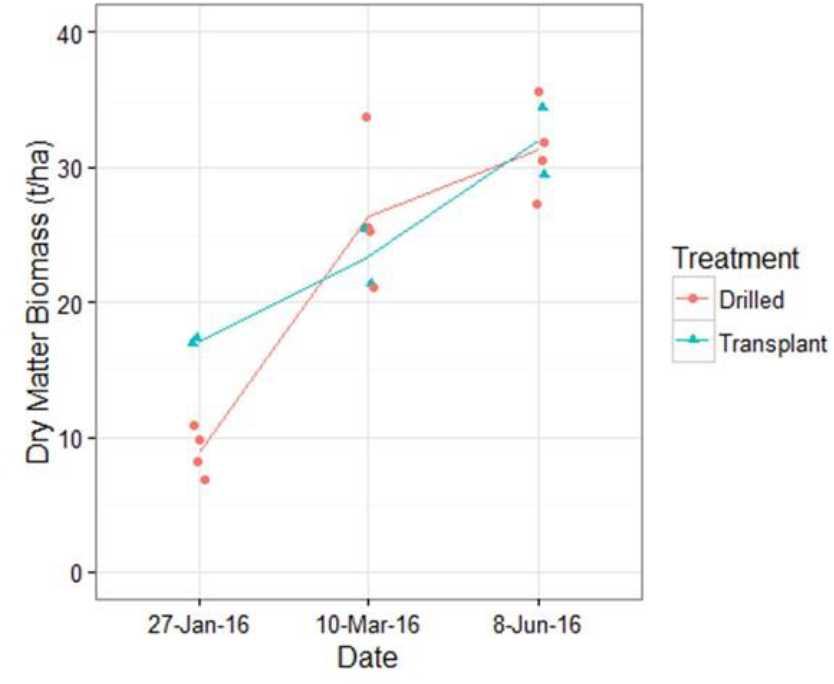
Greater uniformity of the transplanted crop shows here as reduced dispersion. The number of samples from plots at each harvest was two and four from the transplanted and drilled plots, respectively. Of particular note is that the transplanted crop had already reached a common full season yield target by late January indicating potential for early season grazing to fit autumn sowing. The March harvest suggests a similar opportunity for the drilled crop, although the wide dispersion of reported yield in the sampled areas demonstrates increased risk and uncertainty for this strategy.

**Figure 5:**
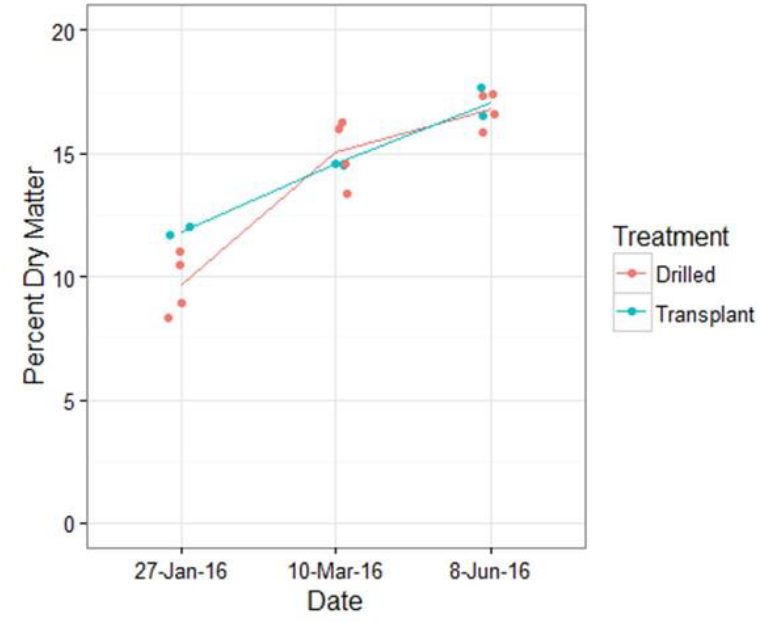
Percentage dry matter in the roots illustrating marginal differences between transplanted and drilled fodder beet crops other than the clear yield advantage offered by transplants early in the season. The number of samples from plots at each harvest was two and four from the transplanted and drilled plots, respectively.

Visual estimates suggested that sprangling of the transplanted crop storage roots might result in differences in root volume between the two establishment methods (Figure 6). In spite of this visual difference, the volumes are essentially the same, except for a greater degree of uniformity of the transplanted crop (Figure 7).

**Figure 6:**
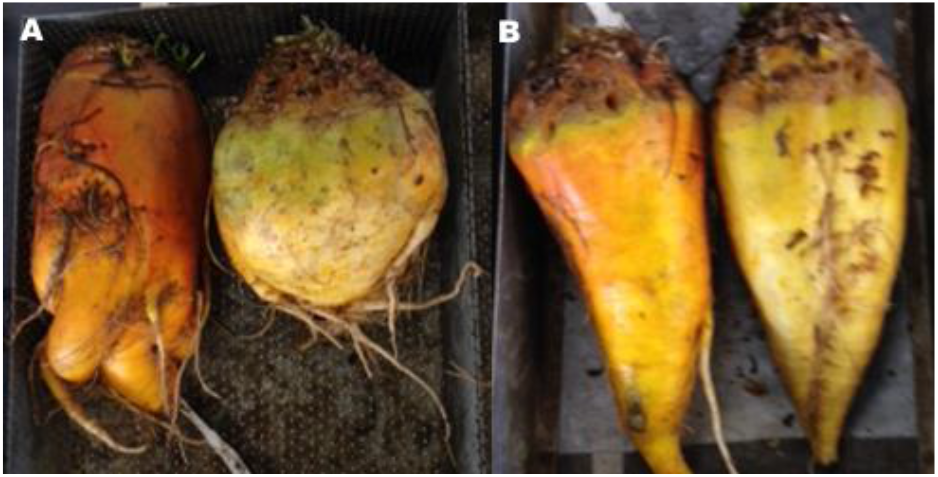
Transplanted (A) and precision-drilled (B) fodder beet storage roots at harvest

**Figure 7:**
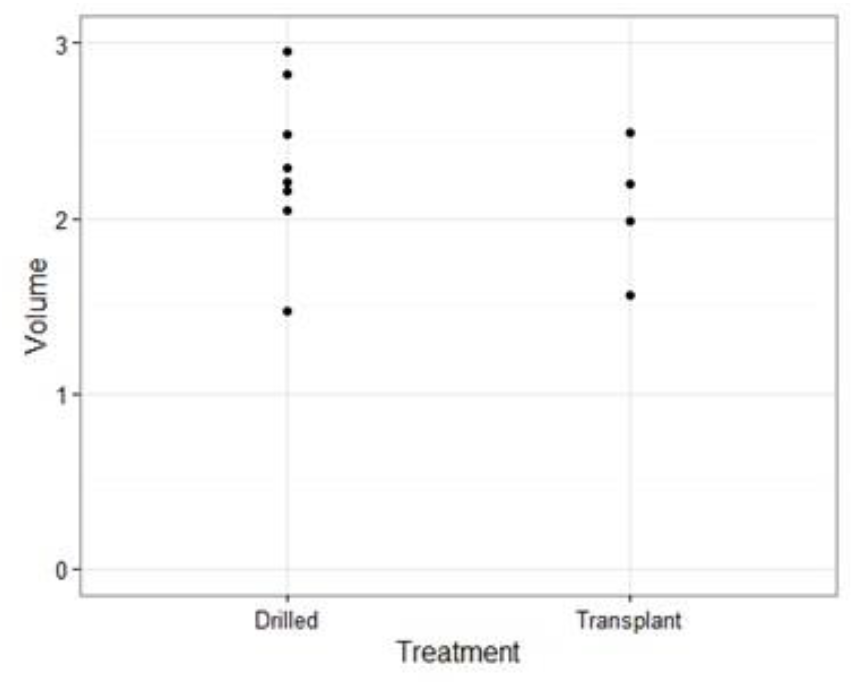
Root volume (litres) at final harvest showing possibly marginal reduction in the transplanted fodder beet crop although this is partially compensated by dry matter content.

Based on our costs of establishment and current price of DM yield, the transplanted crop gave an average gross margin of $2,003/ha, while the drilled crop was $4,971/ha (Table 1).

These results indicate limited difference between transplanted and precision-drilled crops in potential total yield, dry mass percentage and volume of the storage roots. Similar observations have been made for the very closely related transplanted and drilled sugar beet crops in North America (Scott and Bremner 1966; Moraghan and Torkelson 1972; Theurer and Doney 1980). However, the transplanted crop shows a far greater uniformity between plants, and we achieved 100% crop stand. Common experience in the vegetable industry is also near 100% stand establishment. Patchy plant populations in the drilled crops may result in a certain amount of yield compensation, but gaps too large for this compensation effect are most likely the reason for many commercial crops exhibiting relatively poor yields. This is somewhat obscured in this trial as the plant populations for both methods are higher than normal commercial practice. The good stand establishment accounts for the drilled crop yield being considerably higher than commonly reported for commercial crops (Scott and Maley 2010; Milne et al. 2014) and serves as a guide of the true yield potential.

Sprangling of the storage roots (Figure 6) intrinsically suggests a likely yield penalty, but this was not evident in measures of total yield (Figure), DM% (Figure 5), nor, in spite of visual effects, in total volume of the storage roots (Figure 7). This is similar to other experiences from sugar beet trials (Anderson et al. 1958; Moraghan and Torkelson 1972). Different varieties of fodder beet demonstrate different root shape, in particular varying in degree of primary root and hypocotyl proportion forming the storage organ (Milne et al. 2014) and this could result in some differences emerging between the establishment methods.

The potential for an earlier establishment date than typically feasible for drilled crops suggests a distinct yield advantage for transplants where the growing season is limited, either because of cool climatic conditions or necessarily to fit within the farm cropping rotation. In particular, the longer growing season advantage offered by protected cultivation techniques for transplants may be particularly advantageous at higher latitudes (Anderson et al. 1958; Moraghan and Torkelson 1972). The transplants in this trial received some advantage through earlier light interception (Figure 2), although clearly insufficient to derive a total yield advantage. It is likely that transplants could be established considerably earlier than this trial, although further work will be needed to determine the safe date to avoid premature vernalising and bolting (Martin et al. 1983).

Biosecurity is not normally a factor of interest in plant propagation, yet a nursery stage prior to field planting would draw attention to contamination of beet seed with undesirable species. For example, the recent spread of Velvetleaf (*Abutilon theophrasti* Medick) throughout New Zealand via contaminated pelleted beet seed (Ministry for Primary Industries 2016) would not occur via transplanted crops. Firstly, the seedlings look completely different and this should alert nursery managers. Secondly, nursery managers would generally prefer to sow unpelleted seed; velvetleaf and beet seed are visually very different.

Current cost estimates for fodder beet production in New Zealand are $90–$150/tonne DM (Riley 2015) based on 15–25 t DM/ha crops. Our estimates of cost (Table 1) have even our high transplant cost close to the maximum estimate and the drilled crop slightly lower reflecting the unusually high yield achieved under trial conditions. Assuming a “real farm” plant population penalty for the drilled crop, using the same costs but reducing the yield down to 21t DM/ha (Dairy NZ 2016) results in a cost of $137/tonne DM and gross margin of $2371/ha, very close to the transplant crop margin.

There is considerable scope for economising on the costs for transplanting. For example, plant populations as low as 52,000/ha have shown little effect on total yield (Storey et al 1979). Reducing the plant population will offer minimal cost benefit for precision-drilled crops, but will offer very significant savings in a transplanted crop. Some economies are also available by reducing the intensity of field preparation, although a no-till approach is probably not feasible. The cost differential relative to the ease of establishment and security of obtaining a crop from transplants compared to precision-drilling will vary, particularly between growers having to sell the crop as opposed to those growing for their own livestock consumption.

Taking advantage of the earlier season growth available by transplanting may offer significant value in responding to more flexible crop schedules, in particular both crop establishment and grazing earlier than is currently common. For instance the transplanted crops yielded 26 t DM/ha early March which can be grazed to free land for other crops later in autumn. We have not attempted to determine the earliest potential transplanting date and therefore full canopy cover period. This will vary dramatically by latitude, altitude, season and genetic risks of vernalising as a seedling.

## Conclusion

Transplant establishment of fodder beet offers substantial reduction in variation within the crop as well as the potential to take advantage of the nursery phase to extend the effective cropping cycle. Our estimates of cost of production indicate transplanted costs are likely to be about double that of precision-drilled crop establishment, while profit per hectare remains positive. Of particular interest for farmers of this transplant technique for fodder beet is likely to be the security of crop establishment, quality and uniformity of the crop, ease and simplicity involved in establishing a transplanted crop, and thus potential for good yields. Higher latitude crops in particular will benefit from the growing season extension offered by this technique.

## Acknowledgements

Thanks to the Plant & Food, Lincoln field crops research team for growing and harvesting the crop. Plant & Food Research funded the trial.

